# Cross-coronavirus host susceptibility loci influence disease severity through immune mediators

**DOI:** 10.64898/2026.03.23.713300

**Authors:** Ellen L. Risemberg, Sarah R. Leist, Alexandra Schäfer, Kalika D. Kamat, Timothy A. Bell, Pablo Hock, Kara L. Jensen, Colton L. Linnertz, Darla R. Miller, Ginger D. Shaw, Fernando Pardo Manuel de Villena, Ralph S. Baric, Martin T. Ferris, William Valdar

**Affiliations:** Department of Genetics, UNC Chapel Hill; Curriculum in Bioinformatics and Computational Biology, UNC Chapel Hill; Department of Epidemiology, UNC Chapel Hill; Lineberger Comprehensive Cancer Center, UNC Chapel Hill; Department of Microbiology and Immunology, UNC Chapel Hill

## Abstract

Severe disease following infection with SARS-CoV or SARS-CoV-2 is driven in part by genetically regulated immune responses that promote lung injury. Previously, we showed that genetic risk for severe disease is conserved across viruses and between mouse and human, identifying the *HrS43* locus as a shared determinant of severity. Here, to resolve immune pathways linking host loci to disease outcomes, we apply an integrative statistical framework combining Bayesian identification of predictive immune traits with QTL mapping and mediation analysis across infection conditions. This approach identifies immune predictors of disease severity across both viruses, reveals extensive genetic control of the immune system at homeostasis and during infection, and supports locus-specific causal mechanisms of immunopathology. Notably, *HrS43* appears to influence disease severity through distinct immune mediators in SARS-CoV versus SARS-CoV-2, demonstrating that conserved genetic susceptibility can drive virus-specific immunopathology with translational relevance across species.

## 1 Introduction

Zoonotic coronaviruses, including SARS-CoV, MERS-CoV, and SARS-CoV-2, have caused major epidemics with highly variable clinical outcomes. Coronaviruses circulating in animal reservoirs remain poised for human emergence [1] and expanding human-animal contact is expected to increase the risk of zoonotic transmission [2], highlighting the importance of understanding conserved host determinants of disease susceptibility. Human genetic studies have identified multiple loci associated with susceptibility to severe SARS and COVID-19 [3–5], supporting a substantial contribution of host genetic variation to coronavirus infection outcomes. The mechanisms linking these loci to pathological disease responses, however, are often incompletely understood.

Severe coronavirus disease involves substantial immune-mediated pathology [6, 7], particularly during the later phase in which inflammatory lung injury can persist despite low viral burden [8]. Thus, host susceptibility may reflect a predisposition to dysregulated immune responses. Consistent with this, genetic studies have implicated immune pathways such as cytokine signaling and inflammatory cell recruitment in progression of critical COVID-19 [4, 9, 10].

Mouse models enable dissection of genetically driven immune responses by controlling viral strain and dose, timing of infection, and prior immune exposure, while supporting comprehensive immune phenotyping across tissues and disease stages. Genetically diverse mouse populations such as the Collaborative Cross (CC) [11] have enabled mapping of host susceptibility loci for SARS-CoV and SARS-CoV-2 infection [12–16], and have been widely used to characterize genetic regulation of immune variation at homeostasis and in immune-driven disease.

Previously, we used CC-derived populations to demonstrate that some host susceptibility loci are conserved across coronaviruses in mice, including *HrS43*, which is also shared between mouse and human [14–16]. These findings suggest that genetic determinants of coronavirus disease severity may reflect conserved biological pathways relevant to both human infection and future emergent coronaviruses. Here, we integrate immune phenotyping with genetic and disease severity data to investigate immune-mediated mechanisms of genetic susceptibility. We present a statistical framework for jointly modeling correlated immune traits across infection conditions and linking host loci to disease outcomes. Using this framework, we define both virus-specific and shared immune pathways through which conserved genetic susceptibility shapes coronavirus immunopathology.

## 2 Results

### 2.1 CC006×CC044 F_2_ mice exhibit variable disease severity following coronavirus infection

To investigate immune-mediated mechanisms of coronavirus susceptibility, we analyzed disease severity, viral burden, and lung inflammation in a genetic mapping population infected with SARS-CoV MA15 (hereafter, SARS-CoV) [17], SARS-CoV-2 MA10 (hereafter, SARS-CoV-2) [18], or saline (**Fig. 1**). As reported previously, a screen of 10 Collaborative Cross (CC) strains identified CC006/TauUnc (CC006) as susceptible and CC044/Unc (CC044) as resistant to severe disease following infection with SARS-CoV [15]. Despite their divergent disease outcomes, CC006 and CC044 exhibited similar viral loads, suggesting that variation in disease severity reflects differences in host immune responses rather than impaired control of viral replication.

**Fig. 1.**
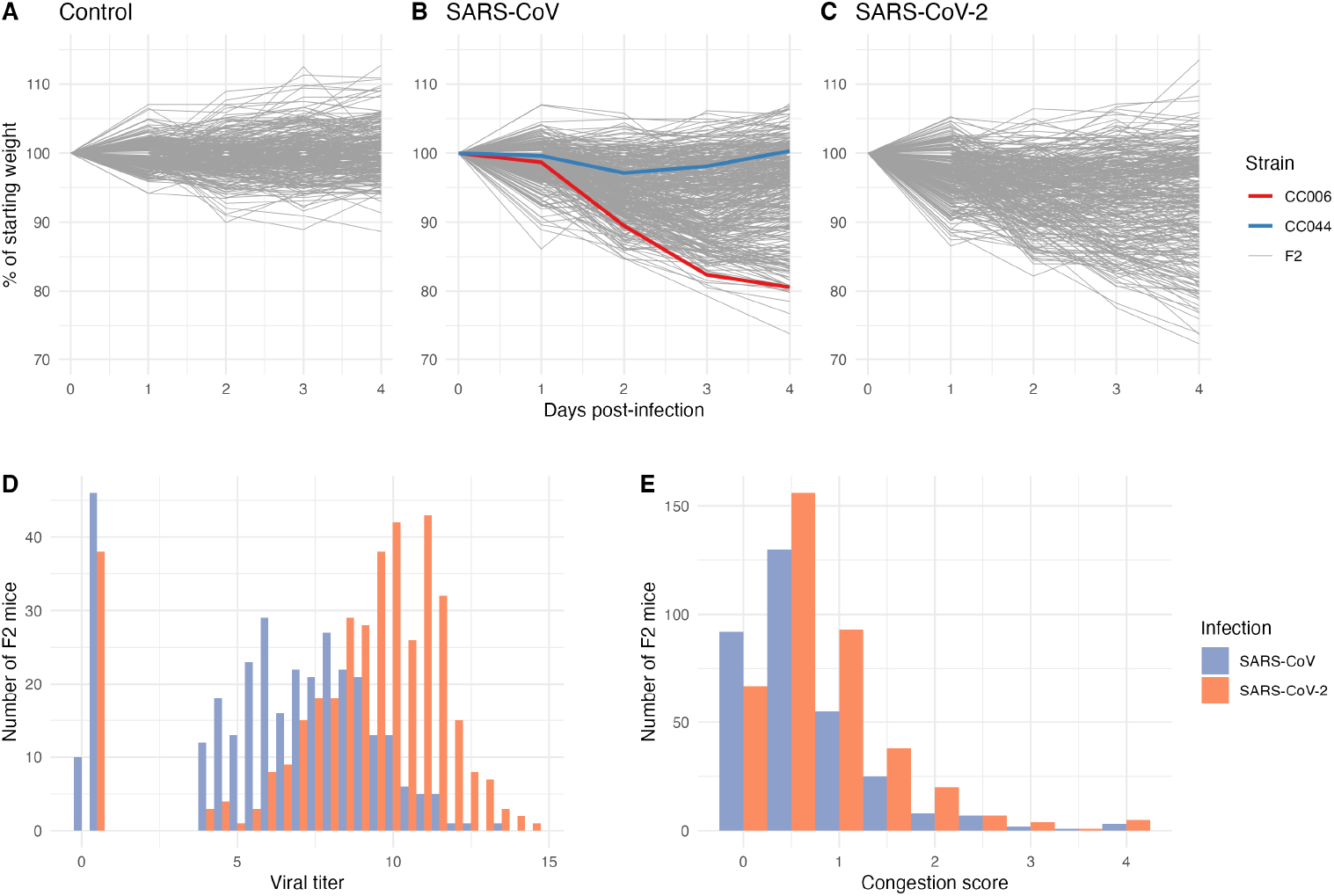
Disease severity traits in F_2_ mice. (A) Weight loss in *n* = 293 F_2_ mice after injection with saline. (B) Weight loss in *n* = 391 F_2_ mice after infection with SARS-CoV MA15, with average weight loss trajectories of parent strains after infection with the same virus. (C) Weight loss in *n* = 324 F_2_ mice after infection with SARS-CoV-2 MA10. (D) Log-transformed viral titer and (E) lung congestion score in infected F_2_ mice, collected from lung tissue at 4 dpi. Weight loss and lung CS in virus-infected mice reported previously [15].

To investigate the genetic and immunological basis of these responses, we generated a CC006×CC044 F_2_ mapping population (hereafter, F_2_ mice) comprising 1008 total mice, each infected with saline (*n* = 293), SARS-CoV (*n* = 391), or SARS-CoV-2 (*n* = 324). Mice were monitored daily for weight loss through 4 days post-infection (dpi) and weight loss trajectories were summarized using an area-above-the-curve (AAC) statistic (**Fig. 1A-C**; **Supp. Fig. A1A**; Online Methods). At 4 dpi, lung tissue was collected to quantify viral titer and pulmonary congestion score (CS) in infected mice (**Fig. 1D,E**). Weight loss and lung CS in infected mice were reported previously [15], and are shown here together with controls and viral burden measurements to establish phenotypic variation in the full mapping cohort.

F_2_ mice exhibited a broad range of disease severity, spanning mild to severe weight loss and lung congestion, as well as substantial variation in viral titers following infection, despite parental strains not differing significantly in viral burden. Viral titers were lower in SARS-CoV-2 than SARS-CoV infection (**Fig. 1D**) and were modestly associated with disease severity (**Supp. Fig. A1B-C**), consistent with contributions of both viral replication and pulmonary inflammation to disease progression.

### 2.2 Lung immune profiling reveals correlated and genetically regulated immune variation

To examine immune mechanisms underlying this variation, we performed flow cytometry on lung tissue from a subset of mice (*n* = 245 control, *n* = 272 SARS-CoV, and *n* = 50 SARS-CoV-2). The smaller SARS-CoV-2 subset of reflects its design for confirmatory rather than exploratory analyses. Using three antibody panels, we quantified 105 immune traits capturing variation in lymphoid and myeloid composition, activation state, antigen presentation, and trafficking receptor expression (Supp. Table 1; Online Methods). Immune traits exhibited modest SNP-based heritability, with several traits showing substantial genetic contribution (**Supp. Fig. A1E**; Supp. Table 1; Online Methods).

Immune traits exhibited extensive multicollinearity, motivating integrative statistical approaches to identify predictive and causal immune traits (**Fig. 2A**; **Supp. Fig. A2**; Online Methods; Supp. Note). Principal components analysis revealed clear clustering by infection group (**Fig. 2B**), and immune profiles distinguished infection status with high accuracy (mean cross-validated accuracy 98.8% ± 0.005%; Online Methods). Together, these data provide a foundation for mapping genetic regulation of immune responses, identifying immune predictors of disease severity, and defining causal pathways that mediate genetic risk.

**Fig. 2.**
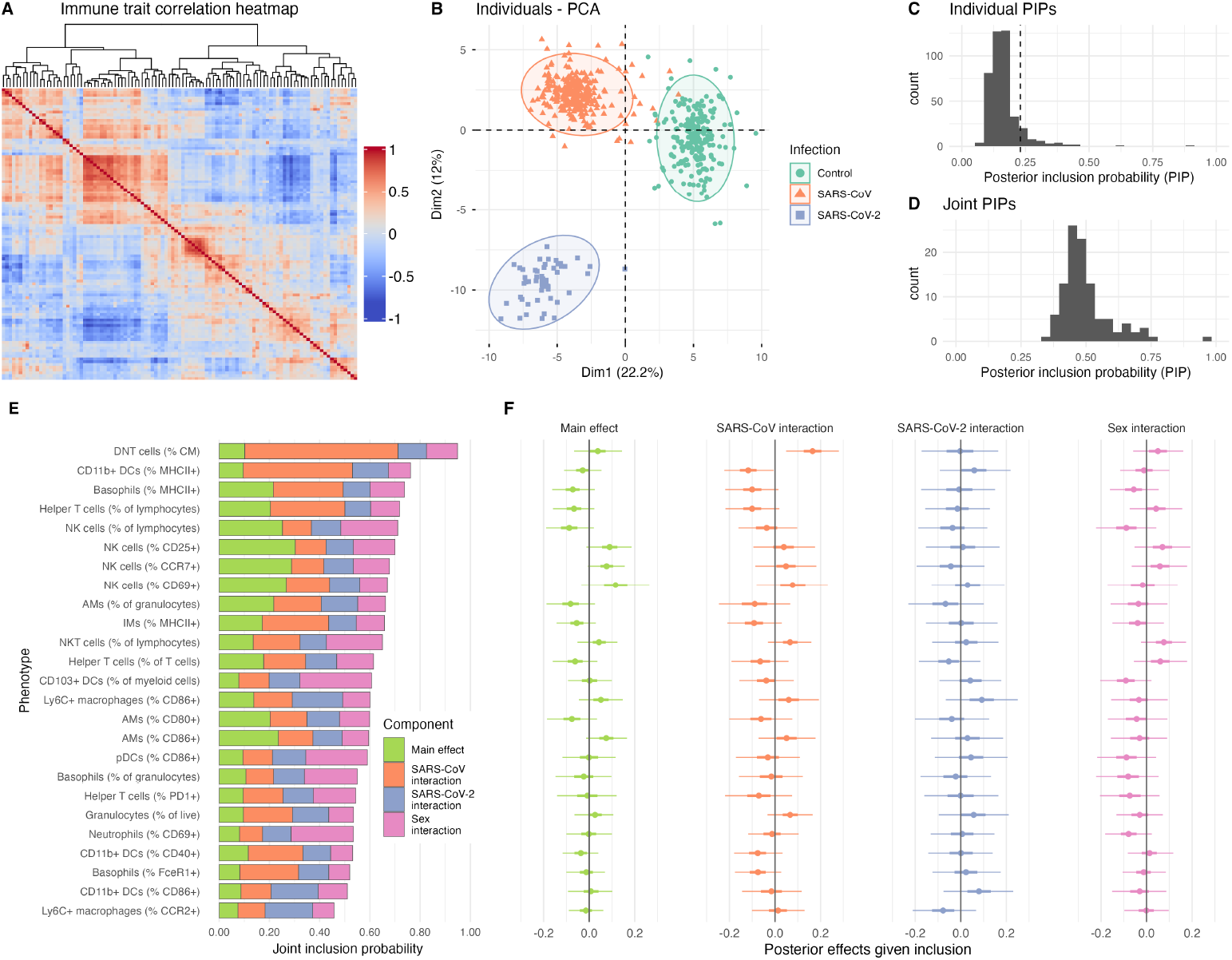
Immune traits and variable selection results. (A) Heatmap of immune trait correlations (labeled version in **Supp. Fig. A2**). (B) PCA of immune traits, colored by infection group. (C) Histogram of individual (per-predictor) and (D) joint posterior inclusion probabilities (PIPs) following Bayesian variable selection. (E) Joint PIPs for selected traits; colors indicate relative importance of main and interaction effects. (F) Posterior effect estimates, conditional on inclusion in the model, with 50% and 95% credible intervals.

### 2.3 Specific immune cell traits are associated with weight loss

To identify immune correlates of disease severity, we applied Bayesian variable selection to model weight loss (AAC) as a function of sex, infection group, and 105 correlated immune traits, which, when allowing for trait-by-sex and trait-by-infection interactions, totaled 420 predictors under selection (Online Methods; Supp. Methods). Variable selection was performed by applying a discrete normal mixture prior to variables under selection (**Fig. A3A**). Our approach accounts for missing immune data by simultaneously performing missing data imputation and variable selection, and accounts for multicollinearity by joint modeling correlated traits and prioritizing traits with high posterior inclusion probability (PIP) and stable selection across cross-validation (CV) folds (**Fig. 2C**; Supp. Note). We summarize evidence for inclusion of each trait using a joint PIP, defined as the posterior probability that any main or interaction effect for a given trait is included (**Fig. 2D**; Supp. Table 1; Online Methods).

The model showed satisfactory convergence (**Fig. A3B**), and explained ∼ 50% of in-sample and ∼ 40% of out-of-sample variance in weight loss, indicating substantial predictive signal in covariates and immune profiles (**Fig. A3C,D**; Online Methods). Thirty-two predictors, representing main and interaction effects from 25 immune traits, exceeded the empirical inclusion threshold and were stably included in CV (**Fig. 2E**; Online Methods). Predictive traits spanned multiple immune compartments, including various subsets of T cells, macrophages, dendritic cells, natural killer (NK) cells and basophils, and describing abundance, activation and differentiation state of these subsets (**Fig. 2E**). Several traits showed largely concordant effects across both viruses, with 19 of 25 traits exhibiting consistent effect direction between SARS-CoV and SARS-CoV-2 infection (Supp. Table 1; Online Methods). For example, helper T cells, alveolar macrophages (AMs) and antigen-presenting (MHCII^+^) interstitial macrophages (IMs) were associated with reduced weight loss in both infections, whereas NK T cells and activated NK subsets were associated with increased weight loss in both infections (**Fig. 2F**). Some traits had infection-specific effects, with, for example, antigen-presenting CD11b^+^ dendritic cells (DCs) having a protective effect after SARS-CoV infection and small deleterious effect after SARS-CoV-2 infection, although this difference was weak and consistent with uncertainty in the SARS-CoV-2 interaction estimate given the smaller SARS-CoV-2 immune cohort (Supp. Note). Together, these results identify a reproducible set of immune predictors of coronavirus disease severity with broadly shared, but occasionally virus-specific, effects.

### 2.4 Immune traits are under genetic regulation

To define genetic determinants of immune variation in the F_2_ mice, we mapped quantitative trait loci (QTL) for 105 lung immune cell traits in infection-stratified and combined analyses. To increase power and explicitly test for context-dependent effects in the combined analysis, we implemented a multi-group mapping framework that jointly models immune traits across infection conditions and explicitly tests for genotype-by-treatment (GxT) interaction effects (Online Methods; Supp. Methods).

We identified 68 significant or suggestive locus-trait associations, corresponding to 41 distinct QTL named *Irq* (Immune response QTL) 1 through 41 (**Fig. 3A**; Supp. Table 2). Eighteen of these loci were only identified in the combined analysis, consisting of small-effect size loci that were underpowered in stratified scans (Supp. Note). Despite the smaller subset of immune phenotyped SARS-CoV-2 infected mice, we detected multiple associations in this group (**Fig. 3A,B**). Comparing effects across conditions revealed both shared and infection-specific architecture: about 38% of locus-trait associations were specific to a single infection group (5 in control, 15 in SARS-CoV, and 6 in SARS-CoV-2), 31% were present in all three conditions, and the remainder were shared across two of the three groups (**Fig. 3B**; Online Methods; Supp. Note).

**Fig. 3.**
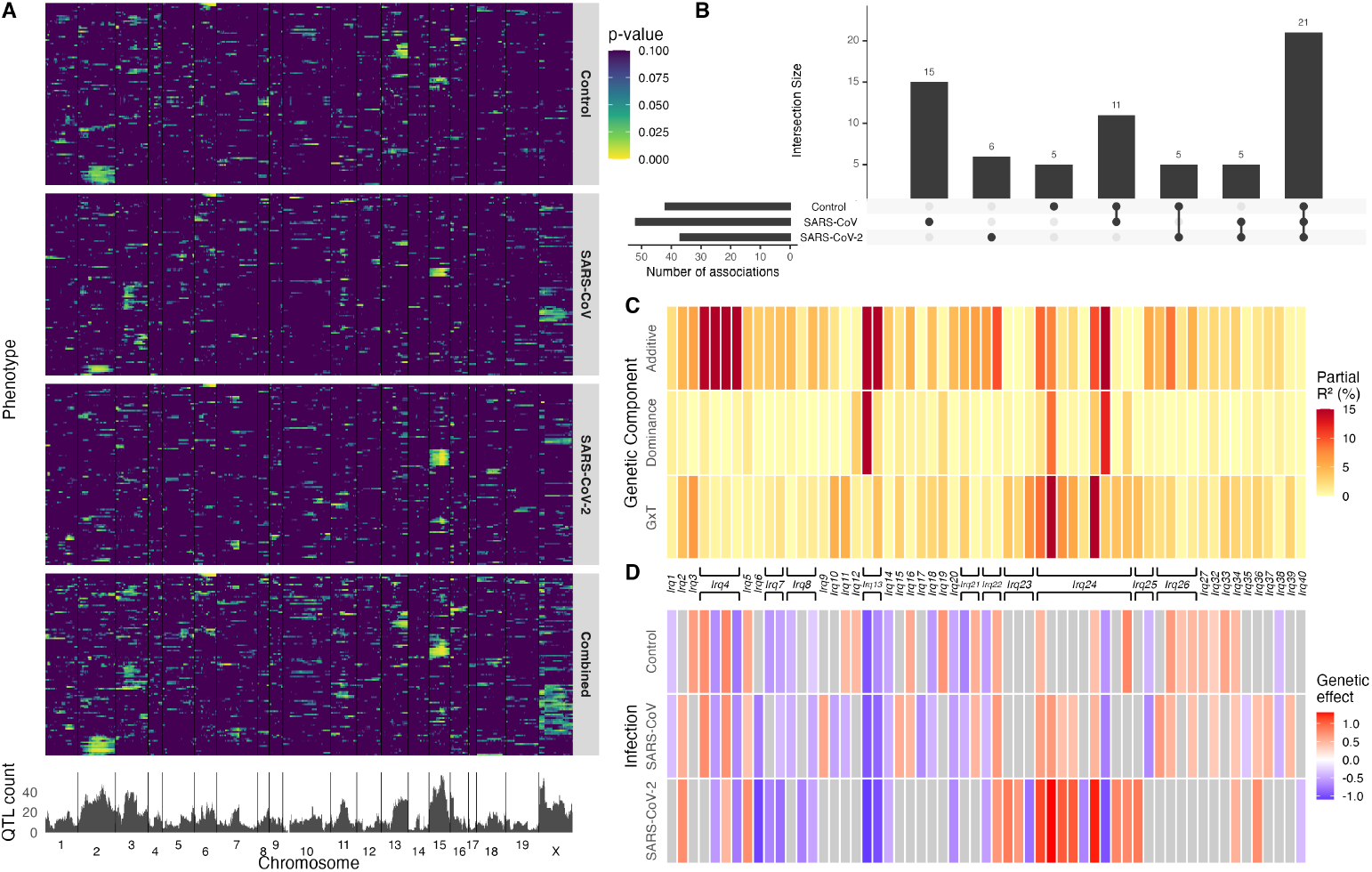
Genetic architecture of immune trait QTL across infection groups. (A) Genome-wide distribution of immune trait QTL across 105 lung immune phenotypes and infection groups, colored by genome-wide adjusted *p*-value. Ordering of traits in this heatmap is specified in Supp. Table 1. (B) Overlap of locus–trait associations (at least nominally significant) across control, SARS-CoV, and SARS-CoV-2 groups. (C) Decomposition of genetic effects at associated loci into additive, dominance, and genotype-by-treatment components. X chromosome QTL are not shown. (D) Genetic effect sizes of immune QTL across infection groups, illustrating shared and context-dependent regulation. Color indicates direction of CC044 allele effect. QTL names and associated traits are provided in Supp. Table 2.

Genetic effects at immune QTL were predominantly additive, with significant additional variance explained by GxT effects and more limited contributions from dominance effects (**Fig. 3C**; Supp. Methods; Supp. Note). Most associations with effects across infection groups were sign concordant (**Fig. 3D**). QTL identified in the SARS-CoV-2 cohort had larger effect sizes than QTL identified in the control or SARS-CoV cohorts (*p* = 1.0 × 10^−4^), reflecting an expected bias towards large-effect QTL in smaller samples (**Fig. 3D**) [19]. We also mapped viral titer using a two-part model [20], given its zero-inflated distribution (**Fig. 1D**). We identified one significant locus in both infection groups, *Irq6*, indicating a genetic contribution to variation in viral burden despite similar titers in parental strains (Supp. Table 2).

### 2.5 Immune traits are mediators of genetically driven disease risk

Having established that immune traits are both associated with disease severity and subject to genetic regulation, we next performed mediation analysis to test whether immune traits mediate genetic disease risk. Mediation analysis is a type of causal inference designed to evaluate whether the effect of an exposure (*X*) on an outcome (*Y*) is explained by a third, mediator variable (*M*), thereby suggesting causal pathways that underly genetic risk. Complete mediation is when *X* acts on *Y* entirely through *M*, and partial mediation is when *X* acts on *Y* only partially through *M* . To perform context-dependent mediation analysis, we extended `bmediatR` [21], a Bayesian model selection approach to mediation analysis, to the multi-group setting. Our approach jointly models control, SARS-CoV, and SARS-CoV-2 cohorts and allows genotype-mediator (*X* → *M*), mediator-outcome (*M* → *Y*) and genotype-outcome (*X* → *Y*) effects to differ by infection group (Online Methods). This framework yields posterior probabilities for complete and partial mediation within each infection group, enabling inference on context-specific mediation patterns within a single probabilistic framework while borrowing information across groups.

We applied this mediation approach to triplets of data consisting of genotypes for immune and disease trait QTL [15], immune traits (including viral titer) as candidate mediators, and weight loss and lung CS as outcomes (Online Methods). Mediation analysis on these triplets evaluates whether the genetic locus affects disease severity through effects on immune variation. We identified 43 locus-immune trait pairs (corresponding to 25 loci and 32 immune traits) with substantial evidence for complete or partial mediation of weight loss or lung congestion (Supp. Table 3). We found that loci with evidence for mediation were often shared across both viruses, although the specific immune trait mediator frequently varied: about half (12) of the 25 genetic loci showed mediation in both infections, but only 4 of these loci had the same mediator across infections.

Evidence for mediated effects at the conserved disease locus *HrS43*, originally reported in Leist (2024) [15], illustrates how shared genetic susceptibility can act through distinct immune mechanisms across infections. The effect of *HrS43* on weight loss was mediated by central memory (CM; CD62L^+^ CD44^+^) double negative (DN) T cells in SARS-CoV infection (**Fig. 4A-D**) but by activated (CD86^+^) inflammatory monocytes in SARS-CoV-2 infection (**Fig. 4E-H**). Specifically, in SARS-CoV-infected mice, the *HrS43* locus is nominally associated with the proportion of DN T cells exhibiting a CM phenotype (**Fig. 4B**), which is in turn associated with increased weight loss (**Fig. 4C**), with strong support for complete mediation (**Fig. 4D**). Whereas, in SARS-CoV-2-infected mice, *HrS43* is associated with the proportion of inflammatory monocytes expressing activation marker CD86 (**Fig. 4F**), which in turn is associated with increased weight loss (**Fig. 4G**), with some evidence for complete mediation (**Fig. 4H**).

**Fig. 4.**
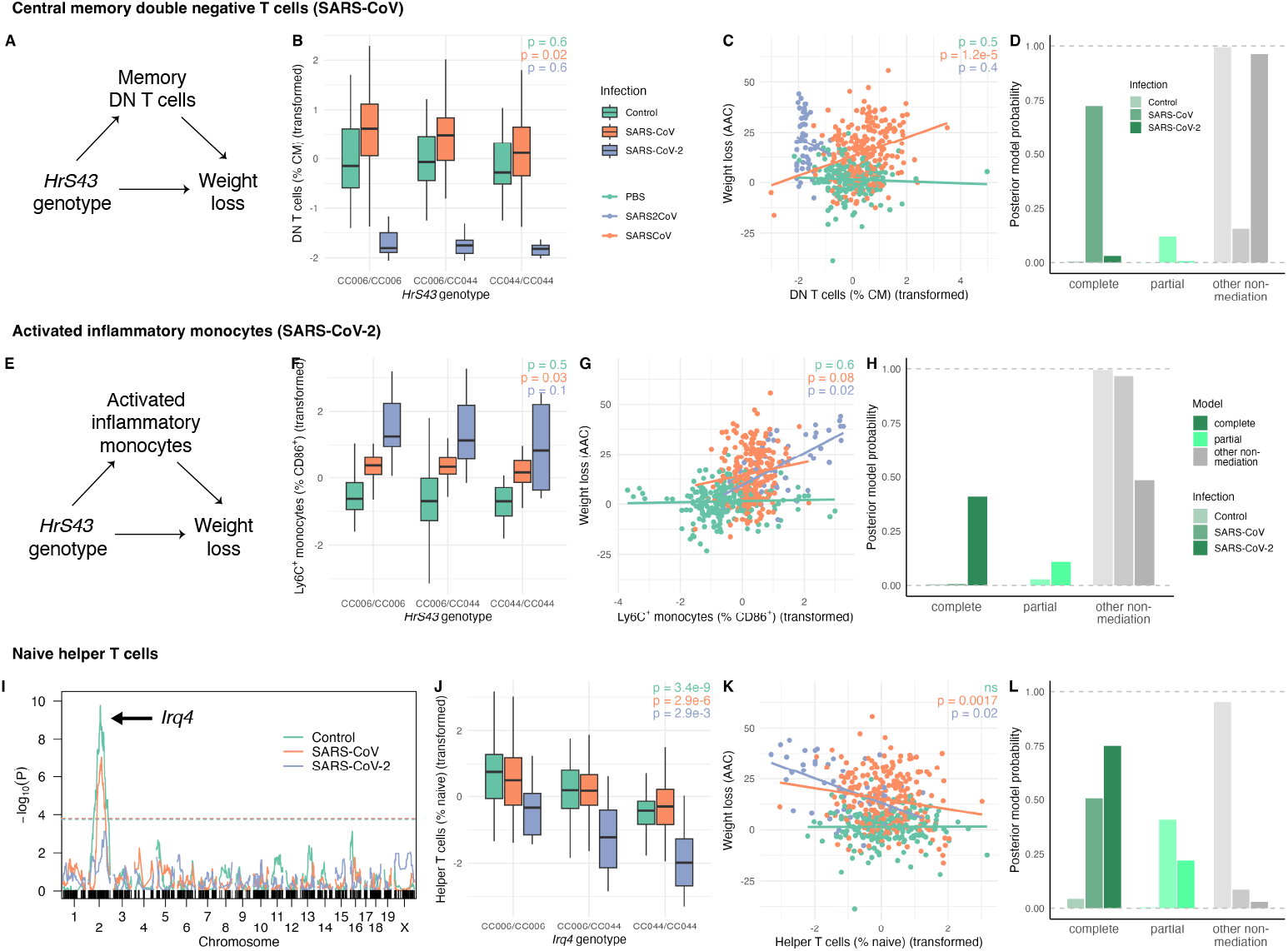
Immune-mediated genetic effects on SARS-CoV and SARS-CoV-2 disease severity. Proportional immune traits were arcsine transformed then centered and scaled prior to analysis and visualization. (A) Schematic of mediation by central memory (CM) double negative (DN) T cells. (B) Effect of *HrS43* genotype on CM DN T cells. (C) Association between CM DN T cells and weight loss. (D) Posterior probability of mediation by CM DN T cells. (E) Schematic of mediation by activated (CD86^+^) inflammatory monocytes. (F) Effect of *HrS43* genotype on activated inflammatory monocytes. (G) Association between activated inflammatory monocytes and weight loss. (H) Posterior probability of mediation by activated inflammatory monocytes. (I) Stratified genome scan of naïve helper T cells. (J) Effect of *Irq4* (chr 2 QTL) on proportion of naïve helper T cells. (K) Association between naïve helper T cells and weight loss. (L) Posterior probability of mediation by naïve helper T cells.

In contrast to *HrS43, Irq4* acts through immune pathways that are conserved across both infections. *Irq4* is associated with the proportion of naïve T cells (**Fig. 4I,J**), which in turn is associated with decreased weight loss (**Fig. 4J**), with strong evidence for mediation in both infection groups (**Fig. 4L**; Supp. Table 3). Increased proportions of naïve T cells were associated with reduced disease severity, consistent with a protective role for preserved naïve T cell compartments during infection. These findings illustrate that, alongside virus-specific immune mechanisms, some genetic loci influence coronavirus disease severity through conserved immunological pathways.

## 3 Discussion

Here we combine Bayesian variable selection, QTL mapping, and context-dependent mediation analysis to dissect genetic regulation of immune traits and their contribution to disease severity. Bayesian variable selection identified a robust set of immune predictors of weight loss with largely consistent effect directions across SARS-CoV and SARS-CoV-2 infection, indicating broadly shared immune protection and pathogenesis. In contrast, QTL mapping of immune traits revealed a mixture of shared and infection-specific genetic architecture, with approximately one third of immune QTL detected in only a single infection group.

Context-dependent mediation analysis further revealed that several QTL have mediated effects on disease severity in both viruses, but the specific immune mediators often vary between viruses. This pattern suggests that host susceptibility loci can influence disease severity across infections while acting through distinct, virus-dependent immune mediators. Consistent with this model, we find evidence that *HrS43*, a disease-associated locus conserved between mouse and human and across the sarbecovirus subgenus [15], acts through different immune traits in each infection: central memory double negative T cells in SARS-CoV, and activated inflammatory monocytes in SARS-CoV-2 (Supp. Table 3). A notable exception to this pattern was *Irq4*, whose effects on disease severity were mediated by naive T cell subsets in both infections, indicating a conserved immune pathway across viruses.

### 3.1 Methodological considerations and inferential scope

Our statistical framework for the integrative analysis of genotypes, immune traits and disease traits has several strengths and limitations. In our Bayesian variable selection analysis, importance is defined in terms of posterior inclusion probability (PIP) and stability across cross-validation folds. Because immune traits measured by flow cytometry are highly correlated, posterior probability mass is often distributed across multiple competing models, reducing the marginal inclusion probability of any single trait despite shared biological signal. Consequently, conservative PIP thresholds may exclude traits that nonetheless contribute to disease severity. Conversely, traits selected by the model should not be interpreted as causal or necessarily of large effect: PIPs quantify evidence that an effect is non-zero, not its magnitude or mechanistic importance. Both selected and non-selected traits should therefore be interpreted in the context of prior biological knowledge.

Mediation analysis also has limitations. Since it requires genotype–immune trait–disease trait triplets, we can only test for locus-specific mediation when genetic regulation of the immune trait is detectable, and only for disease variation attributable to the detected loci. Failure to detect mediation thus does not imply that an immune trait is uninvolved in disease, but rather that mediation by the specified locus could not be established. These constraints are particularly relevant for the SARS-CoV-2 cohort, for which immune profiling was available for a limited subset of animals (*n* = 50), limiting QTL detection. In addition, interpretation relies on assumptions about causal order and direction as well as the absence of unmeasured confounding. Reactive or reverse mediation, where disease severity influences immune composition, is biologically plausible in viral disease but difficult to distinguish statistically [22]. Temporal precedence – that the change in mediator precedes the change in outcome – is also an important condition of causality [23]. The outcome in this study summarizes weight loss trajectories from days 1-4, whereas immune traits were measured on day 4; thus, immune measurements serve as imperfect proxies for immune variation over the course of infection, which may limit strict directional causal interpretation but does not preclude biological relevance.

Nonetheless, because our mediation analysis is based on Bayesian model selection, it offers important advantages. In our framework, evidence for mediation reflects support for a joint model relative to alternative models, rather than requiring each component association to exceed a fixed significance threshold, as in the commonly used “causal steps” approach to mediation [24]. This framework explicitly captures uncertainty in the inferred mediating relationship and allows evaluation of competing causal explanations. For example, CCR2^+^ inflammatory (Ly6C^+^) monocytes were supported as mediators of SARS-CoV-2 disease despite weak marginal associations with weight loss, accompanied by subtantial posterior uncertainty across mediation and non-mediation models (Supp. Table 3).

Additionally, pooling data across infection groups introduces variance heterogeneity that can reduce power and inflate false positives if unmodeled [25]. We address this in both combined QTL mapping and context-dependent mediation analysis using weighted least squares, assigning lower weight to noisier observations. Although unweighted analyses increased detection of QTL and candidate mediators, weighting reduced support for a subset of these signals, suggesting that some associations in unweighted models may reflect spurious associations driven by variance heterogeneity.

Interpretation of SARS-CoV-2-specific results is further shaped by features of the experimental design. Although the overall SARS-CoV-2 infection cohort was similar in size to other groups, immune profiling was performed on a smaller subset of animals, reflecting its design for confirmatory analyses. Nevertheless, several immune QTL were identified in this cohort, with a bias towards large effect size QTL as expected under limited power (**Fig. 3D**). In addition, all SARS-CoV-2 immune profiles were generated in a single experimental batch, resulting in confounding between batch and infection status; however, because F_2_ mice are outbred and littermates were randomized across batches, genetic effects and genotype-by-infection effects remain estimable.

### 3.2 Interpretation of immunological findings

Taken together, the immune traits identified by variable selection and mediation analyses define a focused set of candidates with consistent statistical support for involvement in coronavirus disease pathogenesis. We highlight a small number of representative examples that illustrate two central themes emerging from this study: (i) virus-dependent mediation of shared genetic risk loci, and (ii) convergence on antigen presentation and myeloid subsets that are broadly consistent with observations in human SARS and COVID-19.

A key example of virus-dependent mediation is provided by the conserved disease-associated locus *HrS43*. In SARS-CoV-infected mice, *HrS43* effects on weight loss were mediated by central memory (CM; CD62L^+^ CD44^+^) double negative (DN) T cells. Although canonical memory T cells persist after an infection is resolved, preactivated memory-like DN T cells are often associated with milder respiratory disease [26]. In contrast, the same locus affected SARS-CoV-2 disease severity through activated inflammatory monocytes, a cell type whose accumulation in the lungs has been consistently associated with severe SARS and COVID-19 [27–30].

Beyond this illustrative locus, a broader pattern emerges linking disease protection to antigen presentation and composition of the myeloid compartment. Traits related to macrophages, inflammatory monocytes, and MHCII expression were repeatedly identified as predictors and mediators of disease severity. Increased MHCII expression on dendritic cells (DCs), macrophages and monocyte subsets was associated with reduced disease severity, paralleling human studies in which decreased HLA-DR expression on monocytes and DCs is a hallmark of severe COVID-19 [31–33]. These results support a model in which early preservation of antigen-presenting capacity promotes effective immune coordination and limits immunopathology.

Monocyte/macrophage populations exhibited virus-specific effects. In SARS-CoV infection, genetically driven variation in inflammatory (Ly6C^+^) monocytes mediated disease severity, with inflammatory monocytes having a deleterious effect (Supp. Table 3). This is consistent with evidence in mice and humans that accumulation of inflammatory monocyte/macrophages in the lungs exacerbates SARS-CoV and SARS-CoV-2 disease [27, 28, 30, 34]. Notably, expression of recruitment-mediating chemokine receptor CCR2 on inflammatory monocytes mediated a protective effect on SARS-CoV-2-induced weight loss. Although this appears counterintuitive given the association of inflammatory monocyte recruitment with severe disease in late-stage COVID-19 [4, 35], it is consistent with experimental studies showing that early monocyte recruitment can enhance viral control [36]. This temporal distinction, with prolonged infiltration contributing to pathology, may explain this discrepancy.

Overall, these findings suggest that conserved host susceptibility loci can influence coronavirus disease severity through immune pathways that are often virus-specific, with antigen presentation and myeloid cell abundance emerging as recurrent themes.

By integrating variable selection, genetic mapping, and mediation analysis, this study provides a framework for dissecting how host genetic variation shapes immune mechanisms underlying infectious disease outcomes and highlights immune processes with relevance across mouse models and human coronavirus disease.

## 4 Methods

### 4.1 Experimental methods

#### 4.1.1 Mouse acquisition, production, and usage

Mouse studies were performed in strict accordance with the recommendations in the Guide for the Care and Use of Laboratory Animals of the National Institutes of Health. All mouse studies at UNC (Animal Welfare Assurance #A3410-01) were performed using protocols approved by the UNC Institutional Animal Care and Use Committee (IACUC) in a manner designed to minimize pain and suffering in infected animals.

Collaborative Cross mice were purchased from the Systems Genetics Core Facility at UNC Chapel Hill between 2/2019-11/2020. We generated and maintained sublines of CC006 and CC044 which had identical MHC loci (defined as both having C57BL/6J haplotypes between 34 and 35.5 Mb (b38) on chromosome 17). From these mice, we generated an F_2_ mapping population as previously described ([15]). Briefly, we crossed CC006 and CC044 mice in both directions, to generated reciprocal F_1_s. We then crossed these F_1_s in all 4 possible reciprocal matings to generate F_2_ animals. In this way, potential maternal, mitochondrial, and sex chromosome effects are balanced across our population. At weaning, mice from different birth cages were cohoused together, to ensure that parental or birth-cage effects were distributed across experimental cages. All breeding was conducted in the Department of Comparative Medicine, SPF facilities at UNC Chapel Hill. Mice were given *ad libidum* food and water and kept on a 12:12 light:dark cycle. Experimental mice were transferred into an ABSL-3 laboratory at 9-15 weeks of age, acclimated for one week and then infected. The ABSL3 was kept on a 12:12 light:dark cycle, and mice were given *ad libidum* access to food and water.

#### 4.1.2 Viruses

The generation of recombinant mouse-adapted SARS-CoV MA15 (mouse-adapted SARS-CoV Urbani strain (SARS-CoV-Urbani AY278741) and SARS-CoV-2 MA10 (SARS-CoV-2 Wuhan MN908947) used in this study were previously described ([17, 18, 37]). All experiments were performed in a certified ABSL-3 laboratory, and personnel wore appropriate personal protective gear.

#### 4.1.3 Plaque Assays

Stock viruses and replication competent viral particles in mouse lungs were quantified via plaque assay as reported previously ([17, 18, 37]). Briefly, lung tissue (caudal lobe) was homogenized, and monolayers of Vero E6 cells (ATCC CCL-81) were infected with 10-fold dilutions of supernatant. After incubation for 1 hour at 37°C, infected monolayers were overlayed with agarose in cell media. Resulting plaques were counted after 48 hours (SARS-CoV) or 72 hours (SARS-CoV-2) at 37°C and 5% CO_2_ and final amounts calculated.

#### 4.1.4 Infection

Mice were lightly anesthetized via isoflurane inhalation and infected intranasally with 104 plaque forming units (PFU) of SARS-CoV MA15 and SARS-CoV-2 MA10 in a total of 50µL PBS, or PBS only as mock-infection control. Following infection, mice were assessed daily for weight loss and monitored for mortality. At 4 days post infection (dpi), mice were euthanized via isoflurane overdose followed by bilateral thoracotomy. Mice were perfused with sterile 1xPBS prior to tissue collection. For each animal, the inferior lobe was collected for histological analysis, the caudal lobe for determination of lung titer, the superior lobe for RNA isolation, and samples were stored at -80°C. The entire left inferior lobe was used immediately for the isolation of single cell suspension for flow analysis.

#### 4.1.5 Flow Cytometry staining

We derived three panels of flow cytometry markers to identify granulocyte, myeloid, and lymphoid populations (panel details in Supp. Table 4). Perfused lungs were incubated in digestion buffer (RPMI 1640, 10% fetal bovine serum, 15 mM HEPES, 2.5 mg/ml collagenase A (Worthington), 1.7 mg/ml DNase I (Sigma) for 2 hours at 37°C on a shaker. Cells were then processed through a 70-µm filter, followed by red blood cell lysis with ACK lysis buffer. Single cell suspensions were then diluted to a concentration of approximately 1x107 cells/mL (approx. 1x106 cells/100 µL). 100 µL of each single cell suspension was then stained with a relevant staining panel (Table S1) in staining buffer (1x HBSS, 1% FBS). After staining, the samples were washed 2 times with wash buffer (1xPBS/1% FCS) and resuspended with 200µl antibody mixture (1:200 dilution in staining buffer) of the relevant staining panel described in Table S1 for 60 mins at 37°C. After staining each sample was washed twice (1x HBSS, 1%FBS), and samples were fixed with 2% Formalin, stored in 1xPBS for 24 hrs at 4°C, and then taken out of the BSL3 for analysis.

#### 4.1.6 Flow Cytometry data acquisition and analyses

Following straining, we ran samples on a LSRFortessa (Beckton Dickinson), which is equipped with 5 lasers (355, 405, 488, 561, and 540 nm). Following acquisition, flow cytometry data was analyzed using FlowJo (v10.8.1) to identify a set of 105 immune cell traits, describing the proportional abundance of T cells, B cells, natural killer (NK) cells, macrophages, dendritic cells, and granulocytes (mast cells, basophils, eosinophils, and neutrophils), along with their functional activation states (CD25, CD69, PD1), antigen-presenting status (MHCII), co-stimulatory molecule expression (CD80, CD86, CD40), trafficking receptor expression (CCR2, CCR7), and T cell differentiation state (effector memory, central memory, naïve) (**Supp. Fig. A1D**; Supp. Table 4). Single color compensation controls, fluorescence minus one (FMO) controls and experimental fcs files for lymphoid, myeloid and granulocyte panels were loaded into separate workspaces. A compensation matrix was generated and applied to each panel using the relevant controls. Each sample was quality controlled by removing any artifacts introduced through irregular sample flow or clumping. Single cells were then selected using forward and side scatter height and width. Finally, live cells were chosen by gating on cells excluded by the viability dye (LIVE/DEAD™Fixable Blue Dead Cell Stain Kit, Invitrogen). Each panel was then gated separately to generate lymphocyte, myeloid, and granulocyte phenotypes followed by exporting frequencies of each cell type and mean fluorescence intensities of activation markers.

### 4.2 Statistical methods

#### 4.2.1 Data pre-processing

Immune traits were aggregated from three gated flow cytometry datasets that were gated by two analysts using the same gating scheme. Some observations from the lymphoid panel had multiple measurements, which were harmonized as described in Supp. Methods. All immune traits were logit-transformed and random batch effects were removed (Supp. Methods). Within the subset of mice that were phenotyped for immune traits, some immune trait observations were missing at random due to technical issues with samples.

A statistic representing overall weight loss trajectory for each animal was calculated by measuring the area between a horizontal line at 100% and the line formed by the temperature trajectory in terms of % of starting weight (illustrated in **Supp. Fig. A1A**). This statistic is called area above the curve (AAC).

#### 4.2.2 Heritability estimation

Variance explained by covariates and SNP-based heritability, as described by Yang (2017) [38], were estimated by fitting the linear mixed model,

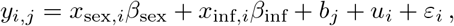

where *y*_*i,j*_ is the phenotypic value for mouse *i* in batch *j, x*_sex,*i*_ and *x*_inf,*i*_ are covariates for sex and infection status and *β*_sex_ and *β*_inf_ are the corresponding fixed effect coefficients, 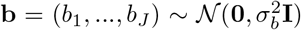 are random batch effects, 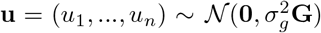 are random polygenic effects based on the genomic relationship matrix **G**, and ***ε*** ∼ 𝒩 (**0**, *σ*^2^**I**) are independent and normally distributed residual errors. This model was fit with `lme4qtl`, and variance components for random effects were extracted from the model using the `VarCorr` function and scaled by total phenotypic variance. Semi-partial *R*^2^ were estimated for fixed effects using the approach described by Nakagawa (2013) [39] and implemented by Jaeger (2017) [40].

#### 4.2.3 Multicollinearity, PCA, and supervised classification

Multicollinearity was evaluated using multiple correlation coefficients and variance inflation factors (VIFs). Traits with VIFs exceeding 5, a common threshold for multi-collinearity [41], were considered multicollinear. Principal components analysis (PCA) was performed on imputed data (using posterior means from the imputation model described below) with R function `prcomp`. We then trained an elastic net multinomial regression classifier to classify infection status from 105 immune phenotypes. We tuned over *α* ∈ [0, 1] and *λ* ∈ [10^−3^, 1], performing 5 repeats of 10-fold cross-validation for each model. The best performing model (*α* = 0.1, *λ* = 0.112) described in Results yielded a mean classification accuracy of 98.8 ± 0.005%.

#### 4.2.4 Bayesian imputation and variable selection

Imputation and variable selection were performed simultaneously so that regression estimates and inclusion probabilities reflected imputation uncertainty. In each iteration of the Gibbs sampler, the missing immune data is imputed in the *imputation phase* and then effects are estimated using non-missing covariates and the current imputed version of the immune data in the *regression phase*.

##### Imputation model

Let **Z** ∈ ℝ^*n*×*q*^ be the matrix containing partially missing immune traits and **X** ∈ ℝ^*n*×*p*^ be the fixed-effect design matrix containing fully observed sex and infection covariates. **Z** follows a matrix normal distribution:

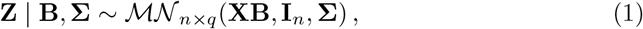

where **B** is the unknown *p* × *q* matrix of regression coefficients, **I**_*n*_ is the *n* × *n* identity matrix, i.e., the rows are independent, and **Σ** is the unknown *q* × *q* residual covariance matrix. Equivalently, each *q*-length row of **Z, z**_*i*_^T^ for *i* ∈ { 1, …, *n* } follows a multivariate normal distribution: **z**_*i*_^T^ | **B, Σ** ∼𝒩_*q*_(**x**_*i*_^T^**B, Σ**).

We use a matrix-normal prior on **B** and an inverse-Wishart prior on **Σ**, with weakly informative prior hyperparameters, resulting in a conjugate matrix-normal posterior on **B** and inverse-Wishart posterior on **Σ** (Supp. Methods) [42]. After updating **B** and **Σ**, missing data in **Z** is imputed for each mouse *i* based on a conditional multivariate normal model as described in Chapter 7.5 of Hoff (2009) [43] (Supp. Methods).

##### Regression model

After imputing missing covariate data, we perform a linear regression with weight loss as a function of fixed covariates, and immune traits that are subject to variable selection. Each immune trait contributes four variables: a main effect, a trait-by-sex effect, a trait-by-SARS-CoV-infection effect, and a trait-by-SARS-CoV-2-infection effect. To identify which of these variables are important in predicting weight loss, we apply Stochastic Search Variable Selection (SSVS) [44, 45] using a discrete normal mixture prior (**Supp. Fig. A3A**). This prior shrinks estimated effects to encourage sparsity and provides posterior inclusion probabilities (PIPs) for each main and interaction effect.

Specifically, let **y** ∈ ℝ^*n*^ be the fully observed weight loss (AAC) phenotype, **X** ∈ ℝ^*n*×*p*^ be the fixed-effect design matrix containing fully observed sex and infection covariates, **Z** ∈ ℝ^*n*×*q*^ be the matrix containing imputed immune traits, and **Z**_int_ ∈ ℝ^*n*×(4×*q*)^ be the matrix containing imputed immune traits and interaction terms with sex and infection, created from the current **Z** in each iteration. Since all covariates in **X** are coded as [− 0.5, 0.5], the resulting **Z**_int_ is still centered and scaled. The multivariate normal likelihood for the outcome is:

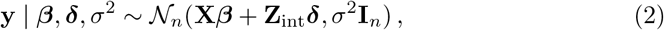

where ***β*** is a *p*-length vector of fixed effects, ***δ*** is a *q*-length vector of estimated coefficients subject to selection, and *σ*^2^ is the residual variance; ***β*** was given a multivariate normal prior, and *σ*^2^ an inverse-gamma prior, both with weakly informative hyper-parameters (see Supp. Methods, including sensitivity analysis). Variable selection was applied to ***δ*** using SSVS [44, 45] with a discrete normal mixture (spike-and-slab) prior, with binary inclusion indicators *γ*_*j*_ ∈ (0, 1) for each *δ*_*j*_, and overall inclusion probability *ψ*,

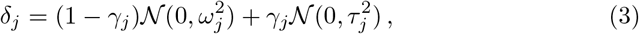

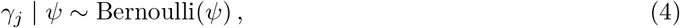

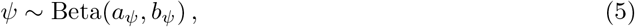

where *τ* ^2^ is the “slab” variance and *ω*^2^ = *τ* ^2^*c*_0_ is the “spike” variance, such that *ω*^2^ ≫ *τ* ^2^. *τ* ^2^ was chosen to be 2.5 × 10^−5^, *c*_0_ to be 300, and prior hyperparameters *a*_*ψ*_ and *b*_*ψ*_ were chosen to center the prior on *ψ* around 0.2. Sensitivity analysis (Supp. Methods) revealed choice of *τ* ^2^, *c*_0_, *a*_*ψ*_, and *b*_*ψ*_ did not meaningfully affect variable selection results.

The posterior Gibbs updates on ***β***, *ψ*, ***γ, δ***, and *σ*^2^ are described in Supp. Methods. The Gibbs sampler was run for 5000 iterations with a burn-in period of 1000 iterations.

##### Posterior inclusion probability thresholds

A posterior inclusion probability (PIP) threshold of 0.5 is common when Bayesian variable selection is performed on orthogonal variables [46], but is considered overly conservative with multicollinear predictor variables [47]. We therefore established a PIP threshold by permutation, permuting the outcome (weight loss) 1,000 times and running the sampler each time using a main-effects only model, thereby establishing a null distribution of PIPs when there is no association between immune phenotypes and disease outcome. The 99th percentile of this null distribution, PIP=0.23, was then used as our selection threshold. (This threshold is conservative, since permutation diagnostics under the interaction model indicated even smaller null PIPs.)

##### Effect size and direction

Posterior means of the virus-specific total effects for each immune trait were calculated as the sum of the posterior means of the main effect and the virus interaction effect, each conditional on inclusion in the model (Supp. Table 1), i.e., for immune trait *j* ∈ [1, *q*] and virus *v* ∈ {1, 2},

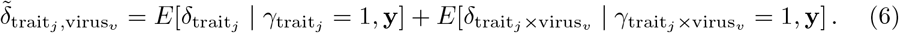

Sign concordance between SARS-CoV and SARS-CoV-2 was established for each immune trait by computing posterior draws of the virus-specific total effect (e.g.,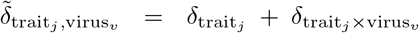) and calculating the posterior probability 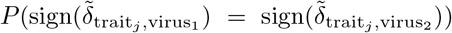. Traits with sign concordance in ≥ 80% of posterior samples were defined as having a “consistent effect direction” between SARS-CoV and SARS-CoV-2.

##### Model performance and stability analysis

Mixing and convergence were evaluated via Gelman and Rubin’s 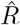 [48]. The portion of variance in **y** explained by the model was estimated using in-sample Bayesian *R*^2^ [49] (**Supp. Fig. A3C**). Overall predictive accuracy was assessed by performing five repeats of 10-fold cross-validation (CV), stratified by sex and infection, estimating the distribution of out-of-sample Bayesian *R*^2^ for each fold, and then aggregating across all folds and repeats (**Supp. Fig. A3D**). The CV results were also used to determine the stability of variable inclusions at our selected PIP threshold by recording whether each variable was selected in each fold and repeat. Stability was defined as the proportion of CV splits in which a variable was selected. Given the high correlation among predictors, we defined stably included variables as those selected in at least 70% of CV folds, balancing reproducibility with the inherent variability in variable selection under correlation (**Supp. Fig. A3E**). We also calculated Jaccard similarity of selected sets of variables across folds at our chosen PIP threshold, which indicates how often the same variables are selected across folds (**Supp. Fig. A3F**).

#### 4.2.5 QTL mapping

Quantitative trait locus (QTL) mapping was performed in two ways: i) stratified by infection group, and ii) a “combined groups” analysis with all three infection groups weighted for heteroscedasticity. In both analyses, we regressed on genotype probabilities as in Haley-Knott regression [50] and handled X-chromosome markers as described by Broman (2006) [51] (Supp. Methods).

In the stratified analysis, the fit of a model with covariates and genotype terms was compared with that of a covariates-only model using an *F* -test, yielding a p-value reported as − log_10_(*p*), hereafter “logP”. In the combined groups analysis, per-marker testing involved comparisons between three (weighted) linear models:

1. Null model: *y* ∼ sex + infection
2. Genotype-only model: *y* ∼ sex + infection + genotype
3. Full model: *y* ∼ sex + infection + genotype + (genotype × infection)

where “infection” is a three-level catagorical variable indicating control, SARS-CoV, or SARS-CoV-2. Model comparisons were conducted using nested *F* -tests to extract three logPs per marker: i) an overall test comparing models (1) and (3), ii) a marginal genotype test comparing models (1) and (2), and iii) a genotype-by-treatment test comparing models (2) and (3).

Pooling infection groups in the combined analysis introduces potentially disruptive variance heterogeneity [25]. To account for this, all models were fit by weighted least squares with group-specific weights. The weights were derived from infection-group–specific residual variance estimates obtained by first fitting the covariate-only linear model

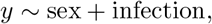

and extracting the residuals. The residuals for each infection group *g* were used to calculate a group-specific variance 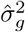, estimated from their sample variance. Observation-level weights were defined as 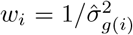, where *g*(*i*) denotes the infection group of individual *i*. These weights were held constant across all markers for a given immune trait.

To assess shared genetic architecture, each locus–trait pair was tested across all groups, and nominal associations (*p* ≤ 0.05) were counted as evidence of shared effects, since testing at pre-identified positions constitutes targeted hypothesis testing rather than genome-wide discovery. QTL effect decomposition into additive, dominance and GxT effects and determination of cross-infection sign concordance are described in Supp. Methods.

##### Multiple testing correction

Genome-wide adjusted *p*-values were determined by permutation test. A standard permutation approach was applied to stratified analyses: for each phenotype, the phenotype was permuted relative to genotype data, a genome scan was performed on the permuted dataset, and the maximum logP value was recorded. This was repeated 1000 times, and the maximum logP’s were used to fit a Generalized Extreme Value distribution, from which genome-wide *p*-values were derived [52, 53]. This permutation test procedure was modified for combined analyses as follows: i) permutations were stratified by sex and infection, ii) weights were recomputed per permutation, and iii) the maximum logP across all markers and all three tests (overall, marginal genetic, genotype-by-treatment) was recorded per scan (as an “omnibus” statistic).

To control for multiple testing across phenotypes, experiment-wide *q*-values were obtained by applying the Benjamini-Hochberg False Discovery Rate (FDR) procedure [54] to the minimum genome-wide adjusted *p*-value for each phenotype within a given group (stratified control, stratified SARS-CoV, stratified SARS-CoV-2, and combined). This provided *q*-values for the top QTL from each scan; for QTL that were not the most significant in a scan, a linear interpolation function mapped sorted *p*-values to their FDR-adjusted *q*-value. QTL were defined as significant if *q* < 0.05 and suggestive if 0.05 ≤ *q* < 0.1.

#### 4.2.6 Mediation analysis

To perform genotype-by-treatment (GxT) mediation analysis, we extended the Bayesian model selection framework of `bmediatR` [21] to the multi-group context. In brief, `bmediatR` performs Bayesian model selection on a set of 2^3^ = 8 causal models defined by the presence or absence of three directed edges, labeled as *a, b*, and *c*, that connect the exposure (*X*), mediator (*M*) and outcome (*Y*) (**Fig. 5A**). Mediation is when edges *a* and *b* are present: complete mediation is when edge *c* is absent, indicating *X* influences *Y* entirely through *M* ; partial mediation is when edge *c* is present, indicating both mediated and direct effects of *X* on *Y* . `bmediatR` also reports colocalization, a specific non-mediation model in which edges *a* and *c* are present, indicating that *X* independently affects both *M* and *Y* . For simplicity, we condense colocalization into the general non-mediation category.

**Fig. 5.**
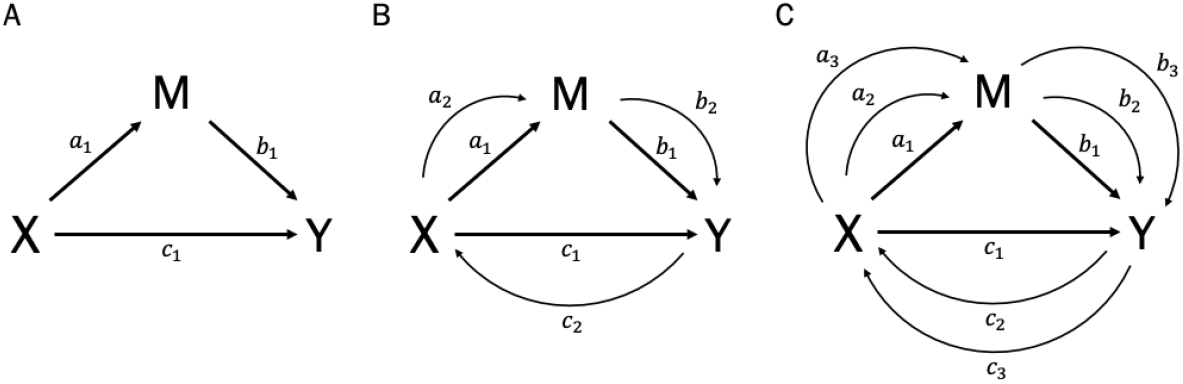
Directed acyclical graphs (DAGs) representing single-group and two-group mediation analysis. DAGs showing edges connecting exposure (*X*), mediator (*M*) and outcome (*y*). In our analyses, *X* represents a QTL genotype, *M* is an immune trait, and *y* is disease severity (weight loss). A) Edge *a* represents the *X* → *M* effect, edge *b* represents the *M* → *y* effect, and *a* and *b* together capture the *indirect* or *mediated* effect. Edge *c* represents the *direct X* → *y* effect. B) Edges *a*_1_ and *a*_2_ represent the *X* → *M* effect in groups 1 and 2, respectively. Edges *b*_1_ and *b*_2_ represent the *M* → *y* effect in groups 1 and 2, respectively. Edges *a*_1_ and *b*_1_ oe *a*_2_ and *b*_2_ capture the mediated effect. Edges *c*_1_ and *c*_2_ represent the direct *X* → *y* effect in groups 1 and 2, respectively.

The framework of `bmediatR` was extended to allow for distinct edges per group (**Fig. 5B,C**) as follows. For *T* groups, group-specific edges are labeled with subscripts to denote group membership, i.e., *a*_*t*_, *b*_*t*_, and *c*_*t*_, and positive evidence for mediation implies that the relevant edges are present within the same group. For example, considering the two-group case, evidence for mediation requires the presence of *a*_1_ and *b*_1_, indicating mediation in group 1; *a*_2_ and *b*_2_, indicating mediation in group 2; or all four, indicating mediation in both groups (with the presence or absence of *c*_1_ and *c*_2_ distinguishing between complete and partial mediation in each group).

The two-group `bmediatR` model results in six edges with 2^6^ = 64 possible causal models (**Fig. 5B**), and the three-group model results in nine edges with 2^9^ = 512 possible causal models (**Fig. 5C**). Each causal model is categorized independently within treatment group as complete mediation (mediated effect only; *a* and *b* present), partial mediation (mediated and direct effects; *a, b*, and *c* present), or non-mediation (any other combination of edges present). Posterior support is aggregated across models to quantify evidence for group-specific mediation and overall mediation.

This multi-group `bmediatR` extension was applied to all QTL identified in this cohort of mice: for immune trait QTL reported in this paper, we test for mediation of weight loss and lung congestion score (CS) by the associated immune trait. For disease-associated loci from Leist (2024) [15] that were nominally associated (*p* < 0.05) with any immune trait, we tested for mediation of the associated disease trait, using the AAC statistic in place of any weight loss trait, by the associated immune trait. The two-group model was used when data was available for only two infection groups (e.g., when the mediator is viral titer and/or the outcome is lung congestion score, which are only available for virus-infected mice); the three-group model when data is available for all three (control and both viruses). To account for variance heterogeneity in pooled data, we use weighted regression, as allowed for in the original `bmediatR` package [21], obtaining weights the same way as in the combined groups QTL mapping. Sex and viral titer (if the mediator itself was not viral titer) were controlled for as fixed effect covariates.

## Acknowledgements

This work was supported by the National Institutes of Health (R35 GM127000 to W.V. and E.L.R, T32 GM135123 to E.L.R, P01 AI181898 to M.T.K., U19 AI100625 to R.S.B and M.T.K., R01 AI157253 to R.S.B and M.T.K.). We thank Mike Love and Mark Heise for their comments on the manuscript. We thank the Systems Genetics Core Facility (UNC) for maintaining and distributing Collaborative Cross mice, the Flow Cytometry Core Facility (UNC) for performing flow cytometry experiments. We thank Jenny Williams and the rest of UNC Research Computing for supporting our high-performance computing needs.

## Data and code availability

The data that support the findings of this study and code to reproduce the results are openly available on GitHub (https://github.com/erisemberg/CoV-immune).

## Appendix A Extended Data

**Fig. A1.**
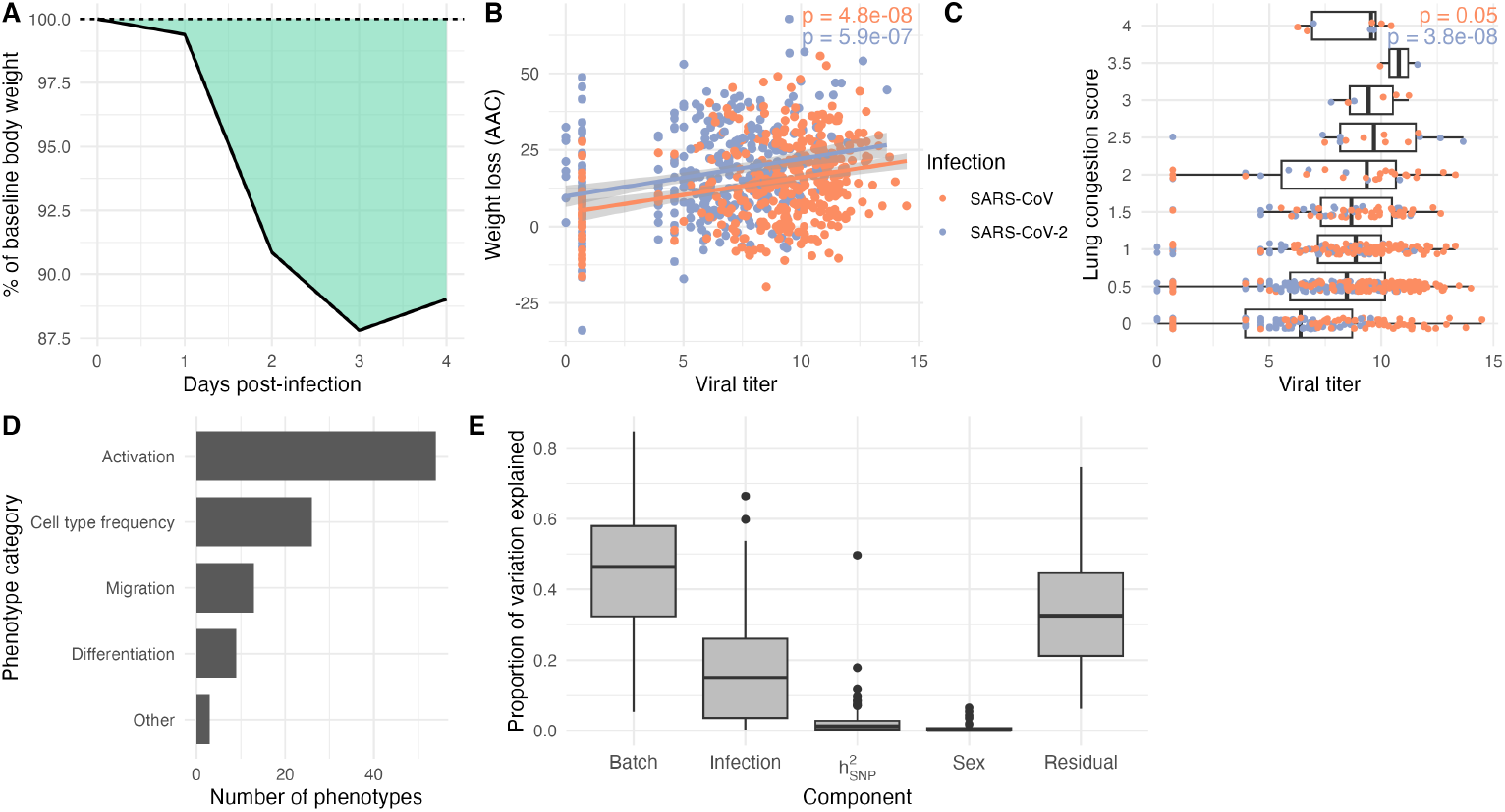
Supplemental trait information. A) Illustration of “area above the curve” (AAC) statistic. B) Association between viral titer in weight loss. C) Association between viral titer and lung congestion score. D) Counts of immune traits by category. E) Variance component analysis.

**Fig. A2.**
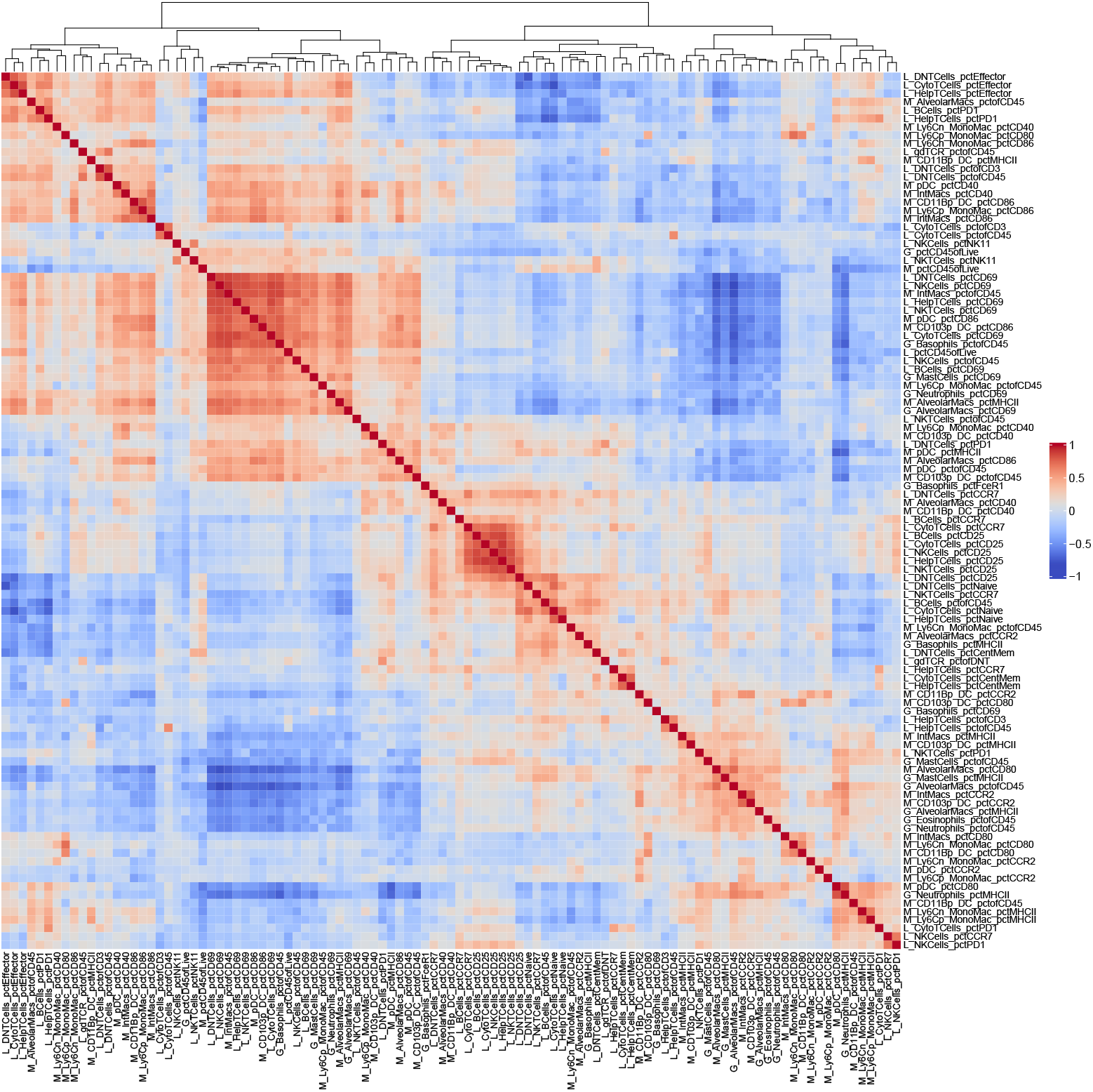
Labeled heatmap of immune trait correlation. Labeled version of heatmap shown in **Fig. 2A**. Hierarchical clustering is based on Euclidian distance of the correlation matrix.

**Fig. A3.**
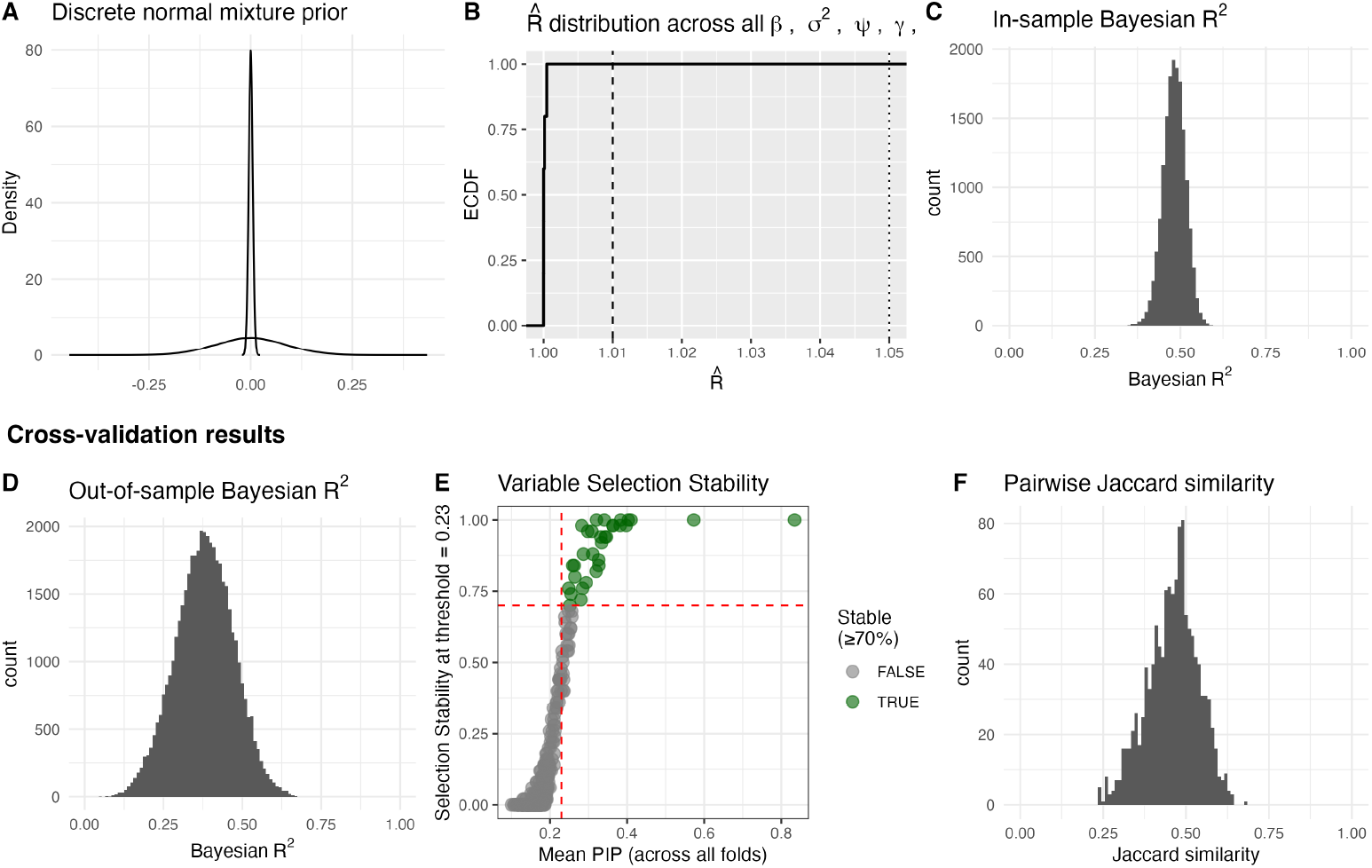
Bayesian variable selection model performance. A) Discrete normal mixture prior defined by chosen hyperparameters *τ* ^2^ = 2.5 × 10^−5^ and *c*_0_ = 300. B) Empirical cumulative distribution function (ECDF) of rank-based Gelman-Rubin statistic 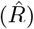 computed across all monitored parameters from multiple independent Gibbs sampling chains. The ECDF at 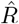 threshold *x* represents the fraction of parameters with 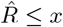. 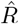 values concentrated near 1 (i.e.,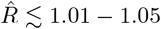) indicate good agreement across chains and are consistent with adequate mixing and convergence. C) Histogram of in-sample Bayesian *R*^2^ values [49] calculated from all post-burn in samples across all chains. D) Histogram of out-of-sample Bayesian *R*^2^ values calculated across 5 repeats of 10-fold cross-validation. E) Mean posterior inclusion probability across CV folds vs. selection stability at our chosen PIP threshold. F) Histogram of pairwise Jaccard similarity of selected sets of variables across CV folds.

