## Supplemental Information for "Cross-coronavirus host susceptibility loci influence disease severity through immune mediators"

#### 1 Supplementary Note

***Section 2.2***

Multiple correlation coefficients ($R^2$) and variance inflation factors (VIFs) were used to assess multicollinearity. Multiple $R^2$s ranged from 0.65 to 0.99 and VIFs exceeded 5, a common threshold for multicollinearity [1], for 90 out of 105 traits. Joint posterior inclusion probabilities (joint PIPs) ranged from 0.36 to 0.95, and from 0.46 to 0.95 for 25 selected traits (**Fig. 2D**).

Posterior inclusion probabilities (PIPs) can be sensitive to hyperparameter choice. To address this concern, we assessed robustness across a range of reasonable hyperparameters and found that, while the mean of the PIP distribution varied, the overall shape and relative ranking of traits within the distribution remained relatively consistent (**Supp. Methods**).

Across important traits, conditional 95% confidence intervals for SARS-CoV-2 interaction effects were larger than those for main, SARS-CoV interaction and sex interaction effects ($p = 3.0 \times 10^{-8}$, Wilcoxon signed rank test; **Fig. 2F**), likely owing to the smaller sample size of immune traits in SARS-CoV-2-infected mice.

***Section 2.3***

QTL identified only by the combined analysis explained less phenotypic variance on average (3.7%) than QTL identified by the stratified approach (19.2%; Welch's two sample $t$-test, $p = 8.1 \times 10^{-8}$). Specifically, pooling data and regressing on both a genotype and GxT term at each marker allows us to detect moderate same-direction effects with the main genetic effect term, and moderate opposite-direction effects with the GxT interaction term. *Irq11*, for example, captures only nominally significant infection-specific effects ($p = 0.03$ in control mice, $p = 0.002$ in SARS-CoV-infected mice) but has a highly significant interaction term ($p = 8.0 \times 10^{-4}$). Since both the stratified and combined analyses identified QTL not detected by the other approach, a combination of approaches is most appropriate in such a treatment-control context.

The increased power to detect QTL in the SARS-CoV infected mice is expected given i) the increased sample size vs SARS-CoV-2 infected mice; and ii) the larger phenotypic variance in infected vs control mice – specifically, 47 of 105 immune traits had significance variance heterogeneity across infection groups ($p < 0.05$, Levene's test), and 39 of these had greater variance in infected mice vs control mice.

Variance decomposition of additive, dominance, and genotype-by-treatment (GxT) effects at each associated locus in each group revealed that genetic effects were predominantly additive, with additive effects explaining 5% of residual variance after accounting for covariates on average (0-44%) (**Fig. 3C**). Three associations were driven by dominance effects – *Irq17*, *Irq19*, and the association of *Irq24* with CCR2$^{+}$ Ly6C$^{+}$ macrophages – highlighting the importance of modeling both additive and dominance effects in genetic mapping (**Fig. 3C**). GxT effects explained a significant amount of variation remaining after accounting for additive and dominance effects (0-26%, mean 3%), supporting the conclusion that many genetic effects are infection-specific or vary across infection groups (**Fig. 3C**).

### 2 Supplementary Methods

##### 2.0.1 Immune trait pre-processing

Flow cytometry data were from three panels: lymphoid, myeloid, and granulocyte and rare DC. All myeloid and granulocyte panels were gated by the same analyst, but lymphoid data was gated by two different analysts: experiments done between 6/20/19 and 1/25/20 were gated by one analyst, with those between 6/20/19 and 11/12/19 being gated twice (by the same analyst), and experiments done between 3/6/20 and 4/15/21 were gated by a second analyst. We integrated the two datasets for experiments between 06/20/19 and 11/12/19 by correcting for an overall effect of the second batch of measurements and then averaging the two values. Thus, if the second measurements are higher or lower than the first measurements on average, we will not introduce that bias into the averaged phenotype. The protocol is as follows:

1. For a given phenotype with two measurements per mouse, let $y_{i,1}$ be the first measurement and $y_{i,2}$ be the second measurement. We fit the following linear model:
$$y_{i,1} \sim \mu + \beta y_{i,2}$$
2. To correct for an overall effect of the second batch of measurements, create a corrected version of the second value as follows:
$$y_{i,2}^{\text{corrected}} = \hat{\mu} + \hat{\beta} y_{i,2}$$
3. Average the first value with the corrected second value:
$$y_i^{\text{avg}} = \text{mean}(y_{i,1}, y_{i,2}^{\text{corrected}})$$

All flow cytometry data were logit-transformed, a common transformation for proportional data that has been shown to eliminate bias in subsequent statistical inference on flow cytometry data [2]. Flow experiment batch effects were handled by fitting the following linear model for all phenotypes,

$$\text{trait} \sim \text{sex} + \text{infection} + (\text{sex} \times \text{infection}) + \text{batch}$$

using R package `lmer` [3], where batch is a random effect. Best linear unbiased predictors (BLUPs) for batch effects were removed from the logit-transformed data (using function `ranef()`), and the resulting data were centered and scaled. In models accounting for both batch effects and gater effects (with batch nested within gater), there were no significant gater effects, so we determined that modeling batch effects was sufficient to account for gater-related variation.

##### 2.0.2 Bayesian imputation and variable selection

We performed imputation and variable selection simultaneously so that regression estimates and inclusion probabilities reflected imputation uncertainty. In each iteration of the Gibbs sampler, the missing immune data is imputed in the *imputation phase* and then effects are estimated using non-missing covariates and the current imputed version of the immune data in the *regression phase.*

***Imputation model***

Missing immune data was imputed using a Matrix-Normal-Inverse-Wishart (MNIW) model as described by Soch (2020) [4]. Let $\mathbf{Z} \in \mathbb{R}^{n\times q}$ be the matrix containing partially missing immune traits and $\mathbf{X} \in \mathbb{R}^{n\times p}$ be the fixed-effect design matrix containing fully observed sex and infection covariates. $\mathbf{Z}$ follows a matrix normal distribution (an extension of the multivariate normal distribution):

$$\mathbf{Z} \mid \mathbf{B}, \boldsymbol{\Sigma} \sim \mathcal{MN}_{n\times q}(\mathbf{XB}, \mathbf{I}_n, \boldsymbol{\Sigma})\,, \tag{1}$$

where $\mathbf{B}$ is the unknown $p \times q$ matrix of regression coefficients, $\mathbf{I}_n$ is the $n \times n$ identity matrix, i.e., the rows are independent, and $\boldsymbol{\Sigma}$ is the unknown $q \times q$ residual covariance matrix. Equivalently, each $q$-length row of $\mathbf{Z}$, ${\mathbf{z}_i}^{\mathrm{T}}$ for $i \in \{1, ..., n\}$ follows a multivariate normal distribution: ${\mathbf{z}_i}^{\mathrm{T}} \mid \mathbf{B}, \boldsymbol{\Sigma} \sim \mathcal{N}_q({\mathbf{x}_i}^{\mathrm{T}}\mathbf{B}, \boldsymbol{\Sigma})$.

We use a matrix-normal prior on $\mathbf{B}$ with $p \times q$ prior mean matrix $\mathbf{M}_0$, $p \times p$ prior row covariance matrix $\boldsymbol{\Lambda}_0$, and $q \times q$ column covariance matrix $\boldsymbol{\Sigma}$ (same $\boldsymbol{\Sigma}$ as above), and an inverse-Wishart prior on $\boldsymbol{\Sigma}$:

$$p(\mathbf{B}) = \mathcal{MN}_{p\times q}(\mathbf{M}_0, \boldsymbol{\Lambda}_0, \boldsymbol{\Sigma}) \tag{2}$$

$$p(\boldsymbol{\Sigma}) = \mathcal{W}^{-1}(\nu_0, \mathbf{S}_0) \tag{3}$$

with weakly informative prior hyperparameters $\mathbf{M}_0 = \mathbf{0}$, $\boldsymbol{\Lambda}_0 = n\mathbf{I}_n$, $\nu_0 = q + 2$, and $\mathbf{S}_0 = (\nu_0 - q - 1)\mathbf{I}_q$. This results in a conjugate matrix-normal posterior on $\mathbf{B}$:

$$\mathbf{B} \mid \boldsymbol{\Sigma} \sim \mathcal{MN}_{p\times q}(\mathbf{M}_n, \boldsymbol{\Lambda}_n, \boldsymbol{\Sigma}), \tag{4}$$

equivalently sampled as: $\mathrm{vec}(\mathbf{B}) \mid \boldsymbol{\Sigma} = \mathcal{N}_{nq}(\mathrm{vec}(\mathbf{M}_n), \boldsymbol{\Sigma} \otimes \boldsymbol{\Lambda}_n)$, where $\boldsymbol{\Lambda}_n = (\boldsymbol{\Lambda}_0^{-1} + \mathbf{X}^{\mathrm{T}}\mathbf{X})^{-1}$ and $\mathbf{M}_n = \boldsymbol{\Lambda}_n(\boldsymbol{\Lambda}_0^{-1}\mathbf{M}_0 + \mathbf{X}^{\mathrm{T}}\mathbf{Z})$; and the residual covariance matrix $\boldsymbol{\Sigma}$ is updated by drawing its precision matrix $\boldsymbol{\Omega}$ from a Wishart distribution:

$$\boldsymbol{\Omega} \sim \mathcal{W}(\nu_n, \mathbf{S}_n^{-1}) \tag{5}$$

where $\nu_n = \nu_0 + n$ and $\mathbf{S}_n = \mathbf{S}_0 + \mathbf{Z}^{\mathrm{T}}\mathbf{Z} + \mathbf{M}_0\mathbf{\Lambda}_0^{-1}\mathbf{M}_0 - \mathbf{M}_n{}^{\mathrm{T}}\mathbf{\Lambda}_n^{-1}\mathbf{M}_n$, and then inverting to recover $\mathbf{\Sigma} = \mathbf{\Omega}^{-1}$.

After updating $\mathbf{B}$ and $\mathbf{\Sigma}$, imputation proceeds based on a conditional multivariate normal model as described in Chapter 7.5 of Hoff (2009) [5]. For each individual $i = 1, ..., n$, let $\mathbf{z}_i{}^{\mathrm{T}}$ be the $q$-length row vector of immune traits (row $i$ of $\mathbf{Z}$). Under the model:

$$\mathbf{z}_i{}^{\mathrm{T}} \mid \mathbf{B}, \mathbf{\Sigma} \sim \mathcal{N}_q(\boldsymbol{\mu}_i{}^{\mathrm{T}}, \mathbf{\Sigma})$$

where $\boldsymbol{\mu}_i{}^{\mathrm{T}} = \mathbf{x}_i{}^{\mathrm{T}}\mathbf{B}$, we partition the observation vector $\mathbf{z}_i$, its mean $\boldsymbol{\mu}_i$, and the covariance matrix $\mathbf{\Sigma}$ according to observed $a$ and missing $b$ immune traits:

$$\mathbf{z}_i = \begin{pmatrix} \mathbf{z}_{i,a} \\ \mathbf{z}_{i,b} \end{pmatrix}, \qquad \boldsymbol{\mu}_i = \begin{pmatrix} \boldsymbol{\mu}_{i,a} \\ \boldsymbol{\mu}_{i,b} \end{pmatrix}, \qquad \mathbf{\Sigma} = \begin{pmatrix} \mathbf{\Sigma}_{aa} & \mathbf{\Sigma}_{ab} \\ \mathbf{\Sigma}_{ba} & \mathbf{\Sigma}_{bb} \end{pmatrix}.$$

We draw the missing data $\mathbf{z}_{i,b}$ from the conditional multivariate normal:

$$\mathbf{z}_{i,b} \mid \mathbf{z}_{i,a}, \mathbf{B}, \mathbf{\Sigma} \sim \mathcal{N}(\boldsymbol{\theta}_{i,b}, \mathbf{\Sigma}_{b|a}),$$

where the conditional mean $\boldsymbol{\theta}_{i,b} = \boldsymbol{\mu}_{i,b} + \mathbf{\Sigma}_{ba}\mathbf{\Sigma}_{aa}^{-1}(\mathbf{z}_{i,a} - \boldsymbol{\mu}_{i,a})$ and the conditional covariance $\mathbf{\Sigma}_{b|a} = \mathbf{\Sigma}_{bb} - \mathbf{\Sigma}_{ba}\mathbf{\Sigma}_{aa}^{-1}\mathbf{\Sigma}_{ab}$.

##### *Regression model*

After imputing missing covariate data, we perform a linear regression with variable selection. Each trait contributed four predictors: a main effect, a trait-by-sex effect, a trait-by-SARS-CoV-infection effect, and a trait-by-SARS-CoV-2-infection effect. We use a discrete normal mixture prior (**Supp. Fig. ??A**) with Stochastic Search Variable Selection (SSVS) [6, 7] applied to immune traits and interaction terms to identify variables important in predicting weight loss. This prior shrinks estimated effects to encourage sparsity and provides posterior inclusion probabilities (PIPs) for each main and interaction effect.

Let $\mathbf{y} \in \mathbb{R}^n$ be the fully observed disease severity phenotype (weight loss as measured by the AAC statistic), $\mathbf{X} \in \mathbb{R}^{n \times p}$ be the fixed-effect design matrix containing fully observed sex and infection covariates, $\mathbf{Z} \in \mathbb{R}^{n \times q}$ be the matrix containing imputed immune traits, and $\mathbf{Z}_{\mathrm{int}} \in \mathbb{R}^{n \times (4 \times q)}$ be the matrix containing imputed immune traits and interaction terms with sex and infection. $\mathbf{Z}_{\mathrm{int}}$ is created from the current $\mathbf{Z}$ in each iteration. Since all covariates in $\mathbf{X}$ are coded as $[-0.5, 0.5]$, the resulting $\mathbf{Z}_{\mathrm{int}}$ is still centered and scaled. The multivariate normal likelihood for the outcome is:

$$\mathbf{y} \mid \boldsymbol{\beta}, \boldsymbol{\delta}, \sigma^2 \sim \mathcal{N}_n(\mathbf{X}\boldsymbol{\beta} + \mathbf{Z}\boldsymbol{\delta}, \sigma^2\mathbf{I}_n), \tag{6}$$

where $\boldsymbol{\beta}$ is a $p$-length vector of estimated fixed effect coefficients, $\boldsymbol{\delta}$ is a $q$-length vector of estimated coefficients subject to selection, and $\sigma^2$ is the residual variance. We apply a multivariate normal prior on the $\boldsymbol{\beta}$ and an inverse-gamma prior on $\sigma^2$:

$$\boldsymbol{\beta} \sim \mathcal{N}_p(\boldsymbol{\mu}_{\beta 0}, \mathbf{I}_p\phi^2) \tag{7}$$

$$\sigma^2 \sim \mathrm{InvGamma}(a_0, b_0). \tag{8}$$

with weakly informative prior hyperparameters $\boldsymbol{\mu}_{\beta 0} = \mathbf{0}$, $\phi^2 = 10 \times \text{svar}(\mathbf{y})$, $a_0 = 3$, $b_0 = (a_0 - 1) \times \text{svar}(\mathbf{y})$, where $\text{svar}(\mathbf{y})$ is the sample variance of the fully observed $\mathbf{y}$ (see sensitivity analysis for details on choice of $a_0$ and $b_0$). SSVS [6, 7] is used to apply variable selection to the predictors in $\mathbf{Z}_{\text{int}}$. This involves a discrete normal mixture (spike-and-slab) prior on the immune effects $\boldsymbol{\delta} = (\delta_1, ..., \delta_q)$, with binary inclusion indicators for each predictor $\gamma_j \in (0, 1)$ and inclusion probability $\psi$:

$$\delta_j = (1 - \gamma_j)\mathcal{N}(0, \omega_j^2) + \gamma_j \mathcal{N}(0, \tau_j^2) \tag{9}$$

$$\gamma_j \mid \psi \sim \text{Bernoulli}(\psi) \tag{10}$$

$$\psi \sim \text{Beta}(a_\psi, b_\psi) \tag{11}$$

where $\tau^2$ is the "spike" variance and $\omega^2 = \tau^2 c_0$ is the "slab" variance, such that $\omega^2 \gg \tau^2$. Hyperparameters were chosen as $\tau^2 = 2.5 \times 10^{-5}$, $c_0 = 300$, with $a_\psi = 2$ and $b_\psi = 7$, which centers the prior $\psi$ around $0.\bar{2}$. Sensitivity analysis revealed choice of $\tau^2, c_0, a_\psi$, and $b_\psi$ did not meaningfully affect variable selection results.

The posterior Gibbs updates on $\boldsymbol{\beta}$, $\psi$, $\boldsymbol{\gamma}$, $\boldsymbol{\delta}$, and $\sigma^2$ proceed as follows.

$$\boldsymbol{\beta} \sim \mathcal{N}_p(\mathbf{A}^{-1}\mathbf{B}, \mathbf{A}^{-1}) \tag{12}$$

where $\mathbf{A} = \mathbf{X}^{\mathrm{T}}\mathbf{X}/\sigma^2 + \mathbf{I}_p\phi^2$ and $\mathbf{B} = \mathbf{X}^{\mathrm{T}}(\mathbf{y} - \mathbf{Z}\boldsymbol{\delta})/\sigma^2 + \mathbf{I}_p\boldsymbol{\mu}_{\beta 0}/\phi^2$.

$$\psi \sim \text{Beta}(a_\psi + \sum \gamma_j, b_\psi + 4q - \sum \gamma_j)\,. \tag{13}$$

Inclusion indicators $\gamma_j$ are updated individually using a collapsed Bayes factor posterior odds update (integrating out $\delta_j$) [8].

1. For each predictor $j$, partial residuals (residuals after removing all terms except the $j$-th term) are computed as $\mathbf{r}_j = \mathbf{y} - \mathbf{X}\boldsymbol{\beta} - \mathbf{Z}\boldsymbol{\delta} + \mathbf{Z}_j\delta_j$, to isolate the evidence for predictor $j$ given the current state of the rest of the model.
2. The collapsed Bayes factor (with $\delta_j$ integrated out), conditional on current $\boldsymbol{\delta}_{-j}$, is calculated by comparing the marginal likelihoods under spike vs. slab priors:

$$\log BF_j = \frac{p(\mathbf{r}_j \mid \gamma_j = 1)}{p(\mathbf{r}_j \mid \gamma_j = 0)} = \frac{1}{2}\left[\left(\log s^2_{\text{slab}} - \log\omega^2\right) - \left(\log s^2_{\text{spike}} - \log\tau^2\right) + \frac{m^2_{\text{slab}}}{s^2_{\text{slab}}} + \frac{m^2_{\text{spike}}}{s^2_{\text{spike}}}\right]$$

   where $s^2_{\text{slab}} = (||\mathbf{Z}_j||^2/\sigma^2 + 1/\omega^2)^{-1}$, $m_{\text{slab}} = s^2_{\text{slab}} \cdot \mathbf{Z}_j{}^{\mathrm{T}} r_j/\sigma^2$, and analogously for the spike prior. The use of $||\mathbf{Z}_j||^2 = \sum_i z^2_{ij}$ enables efficient computation of conditional variances without repeated matrix inversions.
3. The posterior inclusion probability is then:

$$P(\gamma_j = 1 \mid \cdot) = \text{logit}^{-1}(\log\psi - \log(1 - \psi) + \log BF_j),$$

4. and the inclusion indicator for predictor $j$ is drawn from:

$$\gamma_j \sim \text{Bernoulli}(P(\gamma_j = 1 \mid \cdot)) \tag{14}$$

Given current $\boldsymbol{\gamma}$ values, immune effects $\boldsymbol{\delta}$ are drawn from their conditional normal posterior:

$$\boldsymbol{\delta} \mid \cdot \sim \mathcal{N}(\mathbf{A}^{-1}\mathbf{B}, \mathbf{A}^{-1}) \tag{15}$$

where $\mathbf{A} = \mathbf{Z}_{\text{int}}{}^{\text{T}}\mathbf{Z}_{\text{int}}/\sigma^2 + \text{diag}(\mathbf{D}_{\gamma}^{-1})$, $\mathbf{B} = \mathbf{Z}_{\text{int}}{}^{\text{T}}(\mathbf{y} - \mathbf{X}\boldsymbol{\beta})/\sigma^2$, and $\{D_\gamma\}_{j,j} = \omega^2$ if $\gamma_j = 1$ and $\{D_\gamma\}_{j,j} = \tau^2$ if $\gamma_j = 0$. Finally, the residual variance $\sigma^2$ is updated from its inverse-gamma posterior:

$$\sigma^2 \mid \cdot \sim \mathcal{IG}\left(a_0 + \frac{n}{2}, b_0 + \frac{1}{2}(\mathbf{y} - \mathbf{X}\boldsymbol{\beta} - \mathbf{Z}_{\text{int}}\boldsymbol{\delta})^{\text{T}}(\mathbf{y} - \mathbf{X}\boldsymbol{\beta} - \mathbf{Z}_{\text{int}}\boldsymbol{\delta})\right) \tag{16}$$

To prevent premature convergence to an empty model during early learning, two "warm start" strategies are applied for the first 50 iterations: i) the inclusion probability $\psi$ is fixed at its prior mean $a_\psi/(a_\psi + b_\psi)$, and ii) variables identified as top candidates via ordinary least squares regression are held in the model ($\gamma_j = 1$).

***Sensitivity analysis***

We analyzed the sensitivity of the variable selection model to hyperparameter choice by fitting the model over a grid of hyperparameter values: $a_\psi \in (1, 2, 5)$, $b_\psi \in (5, 7, 10)$, $\tau^2 \in (1 \times 10^{-5}, 2.5 \times 10^{-5}, 5 \times 10^{-5})$, $c_0 \in (100, 300, 600)$, and $a_0 \in (3, 5)$ (with $b_0$ calculated from $a_0$ such that the expectation of the inverse-gamma prior on $\sigma^2$, $\frac{b_0}{a_0-1}$, is equal to $\text{svar}(\mathbf{y})$). Leave-one-out cross-validation indicated no statistically meaningful differences in predictive performance, as measured by the expected log pointwise predictive density, ELPD, calculated using Pareto smoothed importance sampling as implemented in the R package `loo` [9], so hyperparameters were chosen for sparsity behavior and interpretability.

##### 2.0.3 QTL mapping

***Genotype probabilities***

For the linear model performed at each marker, genotype probabilities were computed using `R/qtl` [10]. On autosomes, genotype terms consist of additive effects (add $= P(AB) + 2P(BB)$) and dominance effects (dom $= P(AB)$), and on the X-chromosome, genotype probabilities $P(ABf)$, $P(BB)$, and $P(BY)$ are used as genotype terms. Covariates included in all models are sex, infection if performing a combined analysis, and pgm (paternal grandmother) if the marker is on the X-chromosome [11].

For example, the following models were fit at each autosome marker in the combined analysis:

1. Null model: $y \sim \text{sex} + \text{infection}$
2. Genotype-only model: $y \sim \text{sex} + \text{infection} + \text{add} + \text{dom}$
3. Full model: $y \sim \text{sex} + \text{infection} + \text{add} + \text{dom} + (\text{add} \times \text{infection}) + (\text{dom} \times \text{infection})$

Analogous models were fit for X-chromosome markers, with `ABf`, `BB` and `BY` replacing `add` and `dom`, and pgm added as a covariate in all models.

##### ***QTL effect decomposition and sign concordance***

For each autosomal QTL, we fit linear models to decompose genetic effects into additive, dominance, and genotype-by-treatment (GxT) interaction components. Within each infection group, we performed the linear model,

$$y \sim \text{sex} + \text{add} + \text{dom},$$

with `add` and `dom` probabilities defined as described above based on the peak marker. We extracted from this model additive and dominance effect estimates using Type III ANOVA and $p$-values using a sequential Type I ANOVA. QTL were classified as dominance QTL if no infection group showed a suggestive additive effect ($p < 0.1$) but at least one showed a significant dominance effect ($p < 0.05$) after accounting for additivity. The effect size for visualization in **Fig. 3D** was chosen as the additive effect estimate for additive QTL and the dominance effect estimate for dominance QTL. These effect estimates were square-root transformed, preserving sign, and sign concordance across infections was determined by the direction of effects.

To quantify the relative contribution of additive, dominance, and GxT effects across all individuals, we performed the following linear models for each QTL:

$$y \sim \text{sex} + \text{infection} \tag{17}$$

$$y \sim \text{sex} + \text{infection} + \text{add} \tag{18}$$

$$y \sim \text{sex} + \text{infection} + \text{add} + \text{dom} \tag{19}$$

$$y \sim \text{sex} + \text{infection} + \text{add} + \text{dom} + (\text{add} \times \text{infection}) + (\text{dom} \times \text{infection}) \tag{20}$$

and computed partial $R^2$ values sequentially, i.e., the contribution of additive effects was estimated relative to a covariate-only model, of dominance effects relative to the additive model, and of GxT interaction effects relative to the genotype (additive + dominance) model. Partial $R^2$ values were estimated from the residual sum of squares (RSS) for each model as:

$$\frac{\text{RSS}_{\text{reduced}} - \text{RSS}_{\text{full}}}{\text{RSS}_{\text{reduced}}}$$

e.g., the variance explained by dominance effects, as $\frac{\text{RSS}_{18} - \text{RSS}_{19}}{\text{RSS}_{18}}$.
